# Changes in growth, lanthanide binding, and gene expression in *Pseudomonas alloputida* KT2440 in response to light and heavy lanthanides

**DOI:** 10.1101/2024.04.15.589537

**Authors:** Linda Gorniak, Sarah Luise Bucka, Bayan Nasr, Jialan Cao, Steffen Hellmann, Thorsten Schäfer, Martin Westermann, Julia Bechwar, Carl-Eric Wegner

**Affiliations:** Institute of Biodiversity, Aquatic Geomicrobiology, Friedrich Schiller University, Dornburger Str. 159, 07743 Jena, Germany; Department of Physical Chemistry and Microreaction Technology, Institute for Chemistry and Biotechnique, Technische Universität Ilmenau, Weimarerstr. 32, 98693, Ilmenau, Germany; Institute of Geosciences, Applied Geology, Friedrich Schiller University, Burgweg 11, 07749 Jena, Germany; Electron Microscopy Center, Jena University Hospital, Ziegelmühlenweg 1, 07743 Jena, Germany

**Keywords:** Keywords: lanthanides, lanthanome, RNAseq, microfluidics, single-cell ICP-MS

## Abstract

*Pseudomonas alloputida* KT2440 is a ubiquitous, soil-dwelling bacterium that metabolizes recalcitrant and volatile carbon sources. The latter are utilized by two redundant, Ca- and lanthanide (Ln)-dependent, pyrroloquinoline quinone-dependent alcohol dehydrogenases (PQQ ADH), PedE and PedH, whose expression is regulated by Ln availability. *P. alloputida* KT2440 is the best-studied, non-methylotroph in the context of Ln-utilization. We report the most comprehensive differential gene expression analysis, to date, for any Ln-utilizing microbe. Combined with microfluidic cultivation and single-cell elemental analysis, we studied the impact of light and heavy Ln when growing *P. alloputida* KT2440 with 2-phenylethanol as the carbon and energy source. Light Ln (La, Ce, Nd) and a mixture of light and heavy Ln (La, Ce, Nd, Dy, Ho, Er, Yb) had a positive effect on growth, while supplementation with heavy Ln (Dy, Ho, Er, Yb) exerted fitness costs. These were likely a consequence of mismetallation and oxidative stress. Gene expression analysis showed that the Ln sensing and signaling machinery, the two-component system PedS2R2 and PedH, responds differently to (non-)utilizable Ln. We broadened the understanding of the Ln switch in *P. alloputida* KT2440 and could show that it operates as a dimmer switch, modulating the pool of PQQ ADH dependent on Ln availability. Determined quantities of cell-associated Ln suggest a role for Ln beyond alcohol oxidation. The usability of Ln governs the response of *P. alloputida* KT2440 to different Ln elements.

**Importance:** The Ln switch, the inverse regulation of Ca- and Ln-dependent PQQ ADH dependent on Ln availability in organisms featuring both, is central to our understanding of Ln utilization. Although the preference of bacteria for light Ln is well known, the effect of different Ln, light and heavy, on growth and gene expression has rarely been studied. We provide evidence for a dimmer switch-like regulation of Ca- and Ln-dependent PQQ ADH in *P. alloputida* KT2440, and could show that the response to (non-)utilizable Ln differs depending on the element. Ln commonly co-occur in nature. Our findings underline that Ln-utilizing microbes must be able to discriminate between Ln to use them effectively. Considering the prevalence of Ln-dependent proteins in many microbial taxa, more work addressing Ln sensing and signaling is needed. Ln availability likely necessitates different adaptations regarding Ln utilization.

## INTRODUCTION

Together with the chemically similar elements scandium (Sc) and yttrium (Y), the lanthanides (elements 57-71 [La-Lu], Ln) are commonly known as “rare-earth elements” or more precisely “rare-earth metals” (REM). Based on their electron configuration, which governs interactions with other elements, Ln are grouped into light (La-Eu, Sc; LREM) and heavy (Gd-Yb, Y; HREM) REM (1, 2). Ln are key resources for numerous high-tech applications and in increasing demand for the ongoing green energy transition (3). The lack of sustainable mining and recycling methods puts a spotlight on understanding Ln-dependent metabolism as a foundation for biology-inspired Ln recovery strategies.

Our knowledge about Ln-dependent metabolism is primarily derived from methylotrophic bacteria, which utilize C_1_-compounds as carbon and energy sources (4, 5). The first discovered group of Ln-dependent enzymes, Xox-type methanol dehydrogenases (MDH), catalyzes the oxidation of methanol to formaldehyde. The role of Xox-type MDH was long unclear, and calcium-dependent Mxa-type MDH were considered the *de facto* key enzymes for methanol oxidation in methylotrophs (6, 7). Xox- and Mxa-type MDH belong to the diverse group of pyrroloquinoline quinone (PQQ)-dependent alcohol dehydrogenases (ADH) (8). Genes encoding Xox-type MDH were detected in many environments and also in non-canonical methylotrophs (9, 10), suggesting that methylovory (the supplemental use of C_1_ compounds as energy sources) is more common than anticipated (11). The PQQ ADH family includes over a dozen subclades (8). Most are assumed to rely on Ln. In bacteria possessing pairs of Ca- and Ln-dependent ADH, the expression of the corresponding genes is inversely regulated by the “Ln switch”, which is rooted in two-component systems and auxiliary proteins (12–17).

Microbes prefer LREM (5). Mechanisms underlying Ln mobilization and uptake likely differ depending on local Ln bioavailability. Work in the model methylotroph *Methylorubrum extorquens* identified two gene clusters, the *lut*-cluster (lanthanide <u>u</u>tilization and transport) and the *mlu-cluster* (<u>m</u>ethylolanthanin <u>u</u>tilization), which encode uptake machinery centered around a TonB-dependent receptor and an ABC transporter (18), and biosynthetic machinery for a Ln-binding metal chelator (19). Ln commonly co-occur in the environment. It is not clear how Ln-utilizing microbes distinguish between them. Using Beijerinckiaceae bacterium RH AL1, we could show that Ln are selectively taken up and that supplementation with different Ln elements had only minor effects on growth but substantial effects on gene expression (20).

*Pseudomonas alloputida* KT2440 is the best-studied, non-methylotroph in the context of microbial Ln-utilization (15, 21–23). Strain KT2440 is characterized by a broad metabolic versatility, including the potential to use volatile organic compounds (VOC) that carry alcohols or aldehydes as functional groups (24–26). The utilization of VOCs is facilitated by two inversely regulated, broad-range PQQ ADH - PedH (Ln-dependent) and PedE (Ca-dependent). In this study, we investigated the effect of Ln supplementation on growth, cellular Ln, and gene expression. Our findings indicate that the Ln switch operates as a dimmer switch in *P. alloputida* KT2440. Differences in the response to (non-)utilizable elements suggest that *P. alloputida* KT2440 is able to distinguish Ln elements to effectively use them for its metabolism.

## RESULTS

### The effect of La concentration and Ln elements on growth

We first grew *P. alloputida* KT2440 with different La concentrations (10, 50, 100, 250, 500 nM, 1, 2.5, 5, 10 µM; and without added Ln) (**Figure 1A**, left panel). The lag phase was shortest with 10 and 50 nM La. For the corresponding cultures, a maximum growth rate (r) of 1.91 h^-1^ was estimated, almost twice as high as for those without added Ln (**Table S1**). The shortest estimated doubling time (t_d_) values have been 0.36 and 0.38 h in the case of the 10 and 50 nM cultures (**Figure 1B**), respectively. Except for 10 µM, all tested concentrations had a positive effect on r and t_d_ (**Table S1**).

**Figure 1.**
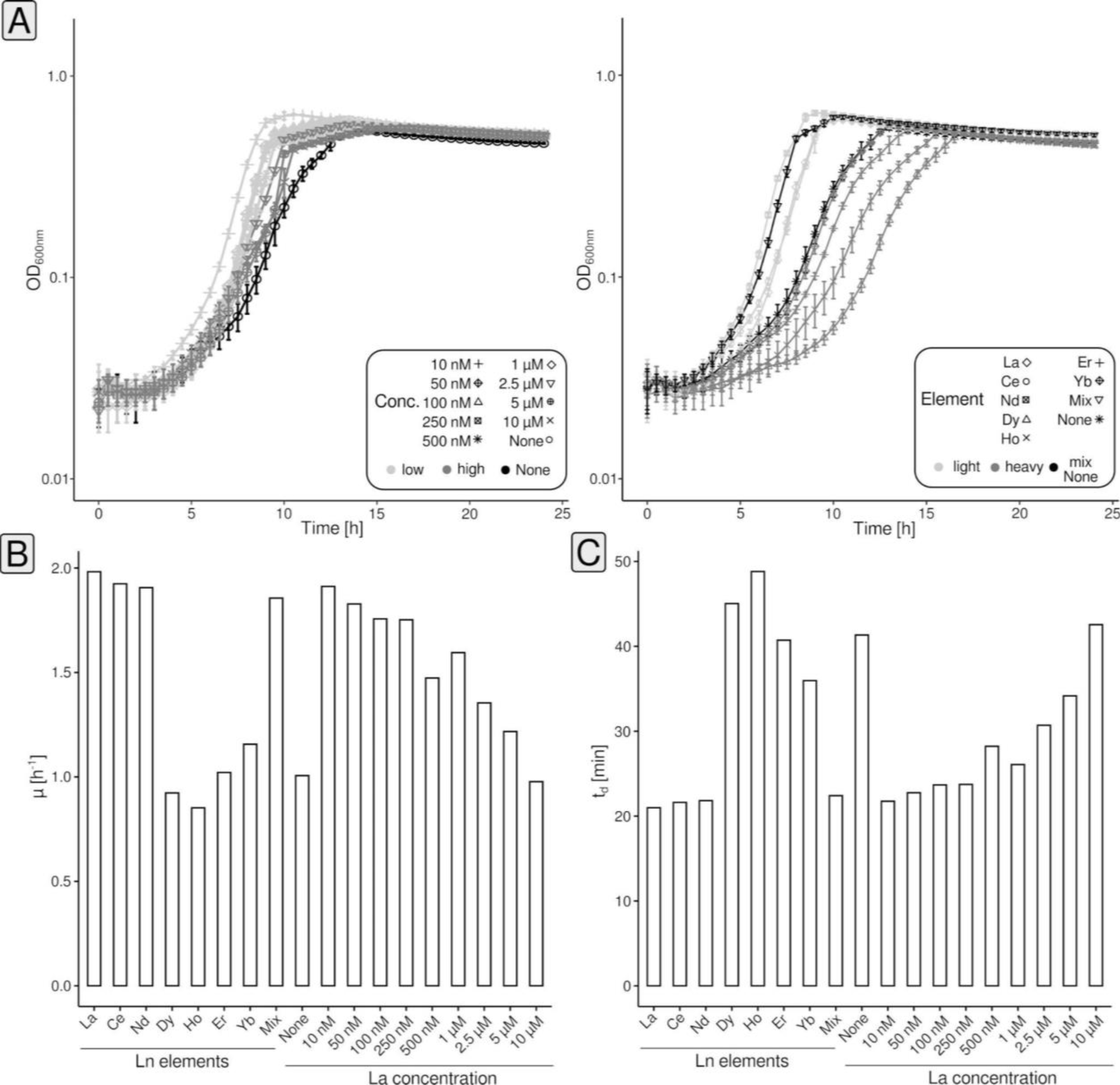
Assessing the growth of *P. alloputida* KT2440 with different concentrations of La and different Ln elements. Cultivations have been carried out in 96-well plates (n = 4 biological replicates) in 300 µL MP medium and growth was tracked through spectrophotometry (OD_600nm_) (A). Growth rates and doubling times were calculated based on averaged data using *growthcurver* (v0.3.1) (74) (B). Obtained growth curve data were fitted based on the standard form of the logistic equation in ecology and evolution, which is rooted in growth rate, initial population size, and carrying capacity (75, 76).

We compared the effect of 50 nM of different Ln elements on growth (La, Ce, Nd, Dy, Ho, Er, Yb, Mix [equimolar mix of Ce, Nd, Dy, Ho, Er, Yb]) (**Figure 1A**, right panel). The lag phase was the shortest for Nd cultures. The lighter Ln (La, Ce, Nd) and the Ln mix had a positive effect on growth when compared to cultures without added Ln (**Figure 1B+C**). Estimated growth rates were between 1.98 (La) and 1.85 h^-1^ (Ln mix), with doubling times around 0.35 h (**Table S1**). The growth rate was 1.00 h^-1^ without added Ln and the doubling time was 0.69 h. Heavier Ln impaired growth. Lag phases and doubling times were longer and growth rates were decreased. Yb cultures almost behaved like those without added Ln.

### Microfluidic one- and two-dimensional screenings with light and heavy Ln

We established microfluidic cultivation (**Figure S1**) using segmented flow for *P. alloputida* KT2440 in 500 nL droplets in PP9 (perfluoromethyldecalin) carrier medium, stored and incubated in PTFE (polytetrafluoroethylene) tubing. The composition concerning added Ln was manipulated yielding either a series of droplets characterized by increasing Ln concentration (0-150 nM) (one-dimensional screenings, **Figure 2A**), or a series of droplets featuring different ratios of Ln concentration (0-75 nM, two-dimensional screenings, **Figure 2B**).

**Figure 2.**
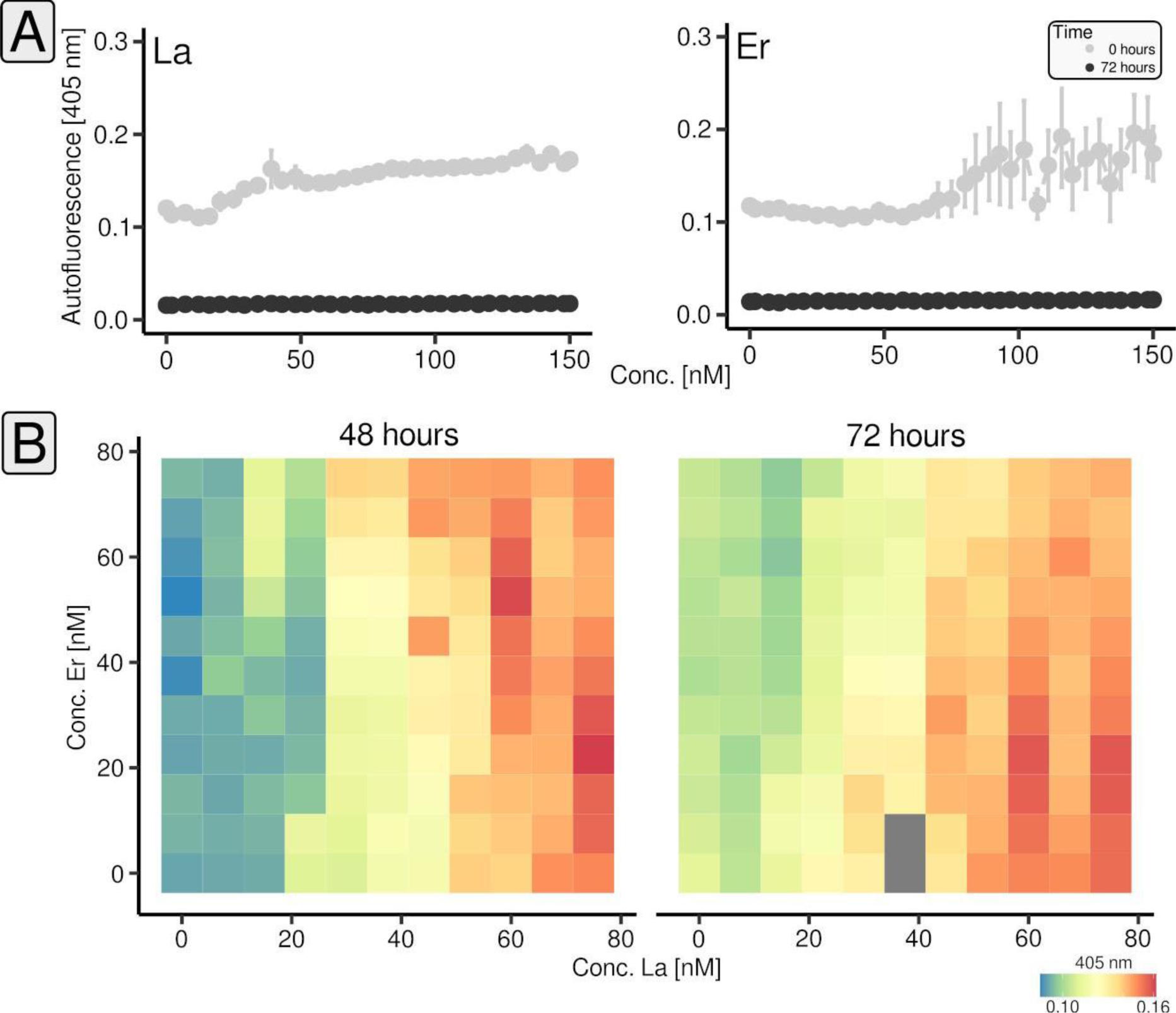
One- and two-dimensional microfluidic cultivation of *P. alloputida* KT2440 with different Ln elements. One-dimensional screenings have been carried out for La and Er ramping from 0-150 nM, over 32 to 35 steps (n = 3-5 microfluidic segments per concentration, technical replicates). Per Ln element, two or three tube coils were incubated (n = 2-3, biological replicates). Incubations were done for 72 hours. Plotted are the average autofluorescence values and standard deviations at the beginning (black) and end of the incubations (grey) (A). A two-dimensional screening has been done for La vs. Er (B). We tested 121 combinations of concentrations (0-75 nM) against each other and incubated the corresponding microfluidic segments for three days. Per combination, between three and five microfluidic segments were generated (technical replicates). Plotted are the average autofluorescence values and standard deviations after two and three days.

*P. alloputida* KT2440 reached the stationary phase in the droplets after 72 hours (data not shown). Growth was monitored by measuring the autofluorescence (405 nm excitation and 425 nm emission). In the case of La, the maximum reached autofluorescence after three days of incubation increased with La concentration, but only marginally for concentrations above 50 nM (**Figure 2A**, left panel). Up to a concentration of approximately 70-75 nM, Er had no impact on reached autofluorescence values, higher concentrations tended to yield increased autofluorescence values (**Figure 2A**, right panel).

We carried out a two-dimensional screening (La vs. Er) to study how the interplay between light and heavy Ln affects growth (**Figure 2B**, **Table S2**). After 48 hours, the highest autofluorescence values were observed for the combinations of 60 nM La and 52.5 nM Er, and 75 nM La and 30 nM Er, respectively (**Figure 2B**). Overall, autofluorescence values tended to be higher with increasing La concentration and if Er was absent or present in lower concentrations. This trend became stronger after 72 hours.

### Analysis of cell-bound Ln using single-cell elemental analysis

The observed effects of different Ln elements on growth prompted us to probe *P. alloputida* KT2440 for Ln accumulation and uptake. Using transmission electron microscopy (TEM) and energy-dispersive X-ray spectroscopy (EDX), we could neither identify extra-nor intracellular Ln accumulations when 50 nM or 1 µM Ln were added (**Figure S2**). To assess cell-associated Ln, we made use of single-cell inductively coupled mass spectrometry (scICP-MS). We measured La, Nd, and Er starting from fixed biomass from incubations supplemented with an excess of Ln (1 µM of the respective element), and measured Ce, Nd, and Er in the case of the Ln mix.

For each sample, several hundred events (= cells) have been analyzed (**Figure 3A**). Cells grown with 1 µM La contained on average 0.058 ± 0.055 fg La × cell^-1^. The cellular Ln content of cells grown with either Nd or Er was higher and around 0.125 ± 0.086 fg Nd × cell^-1^ and 0.152 ± 0.106 fg Er × cell^-1^. Assuming an average wet weight per cell of 2.2 pg (**see materials & methods**), between 2.64 × 10^-3^ (La) and 6.82 × 10^-3^ % (Er) of the wet weight were made up by Ln (**Figure 3B**). For the Ln mix samples, we observed that Ln made up a slightly higher proportion of the wet weight (7.31 × 10^-3^ %) and we could see that the cellular content differed between Ln elements. We detected on average 0.039 ± 0.018 fg Ce × cell^-1^ (1.77 × 10^-3^ % cellular wet weight), 0.059 ± 0.037 fg Nd × cell^-1^ (2.68 × 10^-3^ %), and 0.063 ± 0.014 fg Er × cell^-1^ (2.86 × 10^-3^ %).

**Figure 3.**
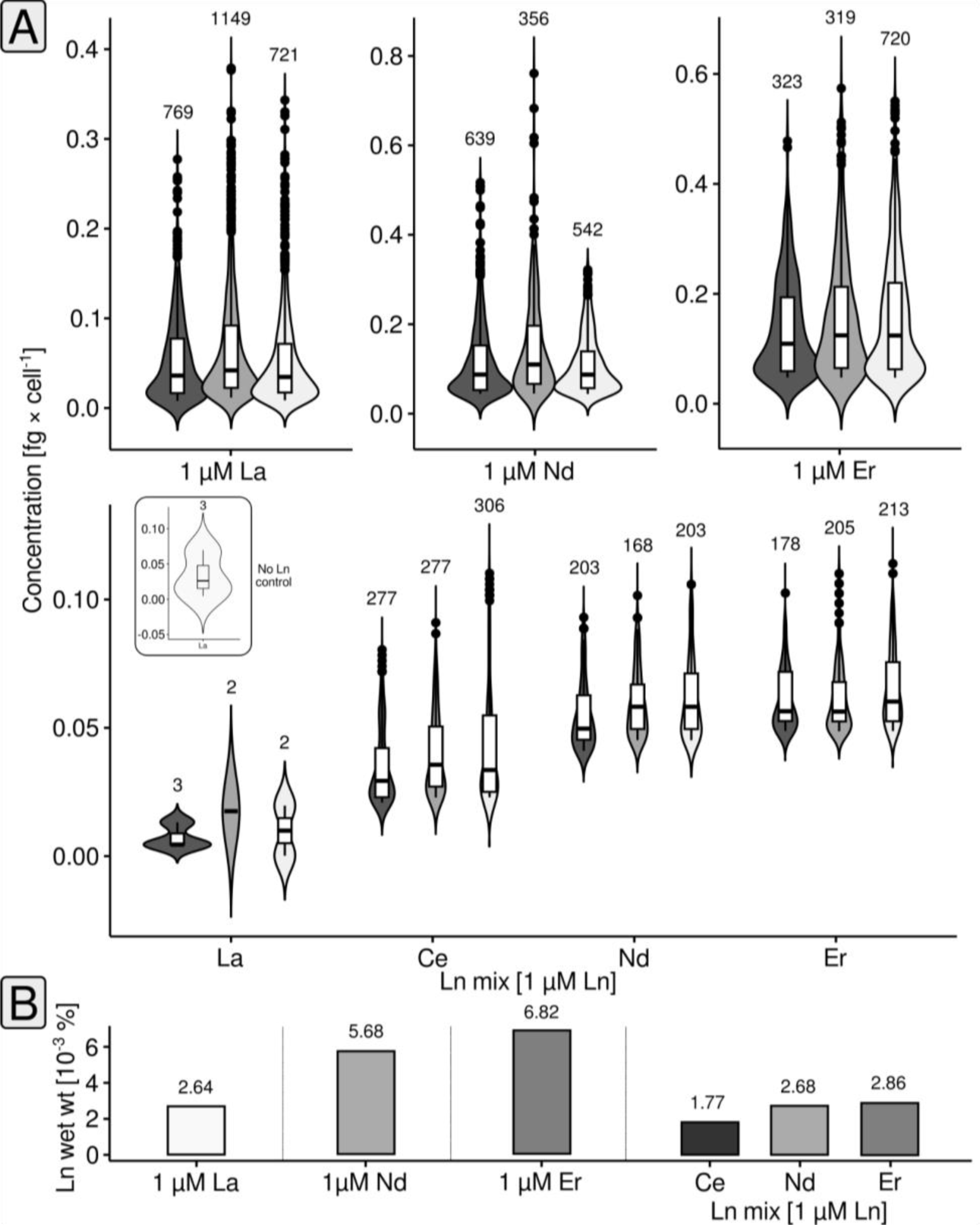
Single-cell elemental analysis. The Ln concentration per cell is shown by combining violin and box plots (A). The box plots indicate lower and upper quartiles, as well as median values. Black circles represent cells with concentrations above (or below) the whisker limits. The numbers above the violin plots indicate the number of considered events (= cells). The small subpanel shows the results of the control sample without added Ln. In this case, we detected three La events, which represent background noise of the measurement. The % Ln of wet weight (B) was calculated as outlined in the materials and methods.

### Overall effects of Ln elements on gene expression

Starting from mid-exponential phase biomass samples (t_1_ + t_2_, **Figure S3**), we performed RNAseq-based (**Table S3**) differential gene expression analysis (DGEA). We considered genes with changes in the expression above |0.58| log_2_FC (fold change; a log_2_FC > |0.58| is equivalent to gene expression changes > 50%), expression values higher than 4 log_2_CPM (counts per million), and false-discovery rate (FDR)-adjusted p-values < 0.05 as differentially expressed (**Figure 4A**, **Table S4-13**). We determined differentially expressed genes (DEGs) for a total of 10 different comparisons: (1) La vs. Er, (2) Nd vs. Er, (3) Mix vs. Er, (4) La vs. None, (5) Nd vs. none, (6) Mix vs. none, (7) Nd vs. La, (8) Mix vs. La, (9) Mix vs. Nd, and (10) Er vs. None (**Figure 4A**).

**Figure 4.**
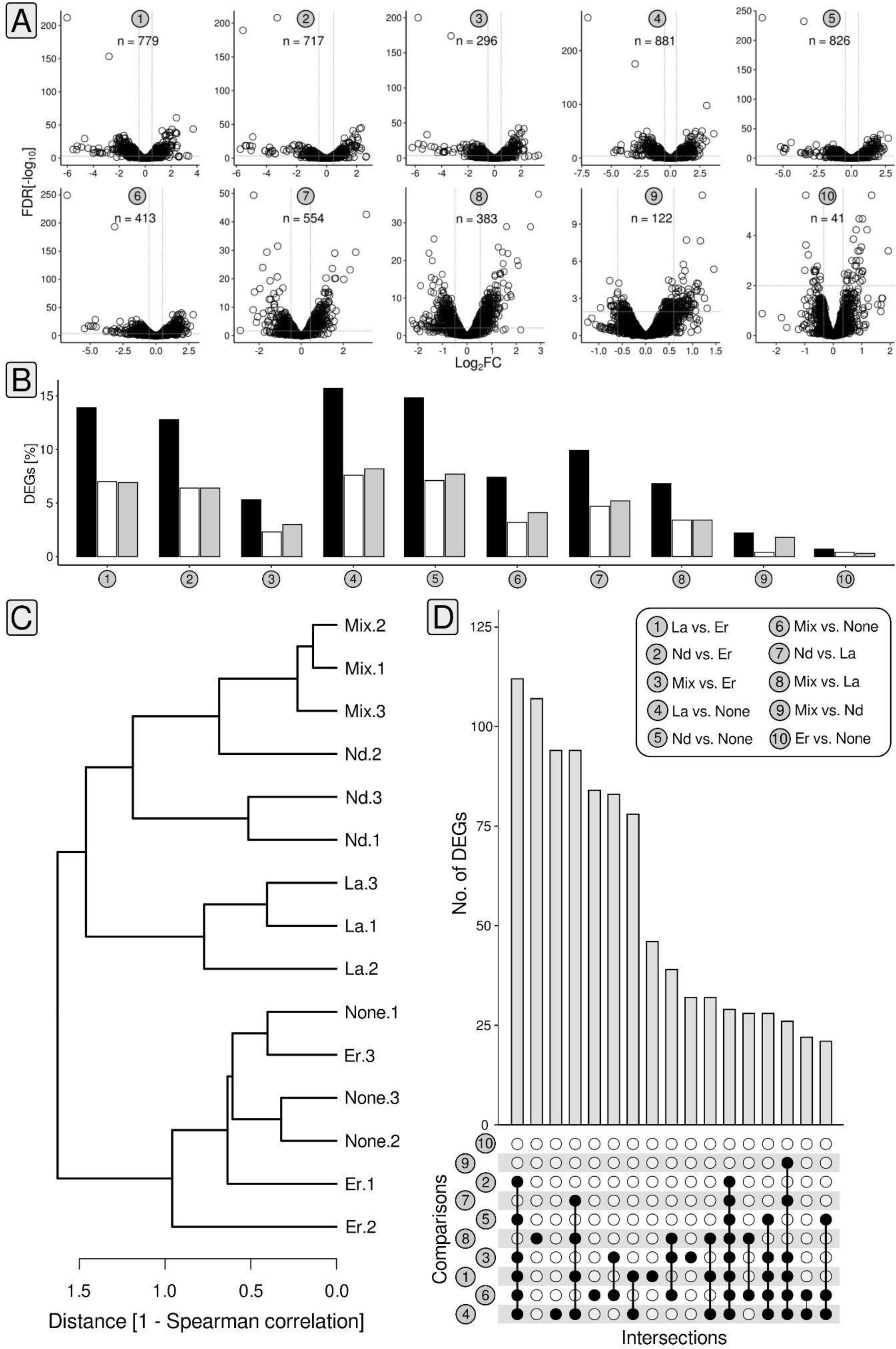
Statistics and summary of gene expression analysis. Volcano plots summarize the number of differentially expressed genes (DEGs) and the breadth of observed fold changes for the different comparisons (1)-(10). We defined differential gene expression based on changes in expression (> |0.58| Log_2_FC [fold change]), relative gene expression (> 4 log_2_CPM [counts per million]), and FDR-corrected p-values (< 0.5). Grey dashed lines highlight the log_2_FC threshold of |0.58| (A). The proportion of DEGs was summarized as bar charts, including the proportion of up- and downregulated genes (B). Based on the counts per million (CPM) matrix, inter-sample relationships have been determined by calculating Spearman distances (C). Shared genes between the different comparisons were highlighted using UpSet plots (D).

The number of DEGs ranged between 41 (Er vs. None) and 881 (La vs. None), which translates into 0.7 and 15.7 % of the *P. alloputida* KT2440 protein-encoding genes (**Figure 4B**). Replicate data sets from the incubations with Er and without added Ln clustered together. The same was true for the La, Nd, and Ln mix replicates (**Figure 4C**).

We determined the overlap between individual comparisons (**Figure 4D**) and noted the biggest overlap of 112 DEGs when looking at comparisons (1)-(6). 94 shared DEGs were observed in the case of (1), (4), (7), and (8). 28 genes were differentially expressed in all comparisons except (9) and (10). There were no DGEs that were shared by all comparisons.

### Differential gene expression and cluster analysis

We used *K-means* clustering (27) for grouping differentially expressed genes based on co-expression. We evaluated different algorithms (**Figure S4**) for estimating K, and ultimately settled on K = 6. Inspecting gene expression by determined z-scores, revealed pairs of clusters characterized by inversed gene expression patterns (clusters 1 and 3, 2 and 4, 5 and 6) (**Figure 5A**).

**Figure 5.**
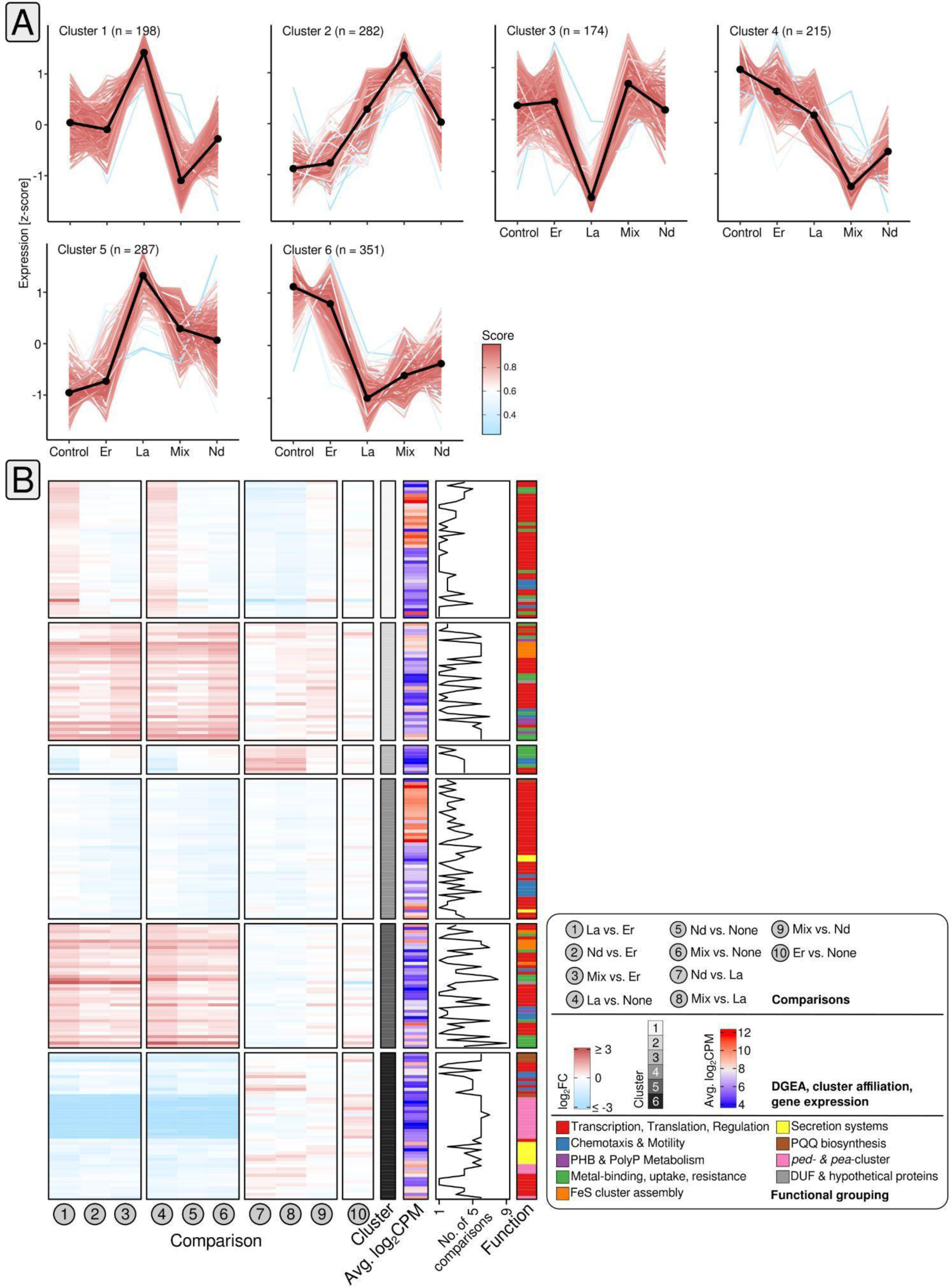
Cluster and differential gene expression analysis. K-means clustering (27) identifies clusters of genes with matching gene expression patterns (A). The number of clusters was estimated *a priori* through Spearman distances, the Calinski-Harabasz index (77), the sum of squared errors, the average silhouette width (78), and the gap statistic (79) (**Figure S4**). We ultimately settled on K=6. The color code indicates the Pearson correlation between each gene in the respective cluster with the cluster centroid. Changes in gene expression of cluster genes for each comparison ((1) - (10)) are shown for selected subsets of genes (B). Heatmap color codes refer to log_2_FC and average log_2_CPM values. The functional annotation of the genes is color-coded as well. The line plot indicates how often the genes were considered as differentially expressed based on the different comparisons.

We noted that cluster 1 genes responded to La supplementation by increased gene expression compared to Er and cultures without Ln, but also reduced gene expression when comparing La to Nd and the Ln mix. Cluster 1 included ribosomal protein (PP_0445-46) and transcriptional regulator (PP_4476) genes (**Figure 5B**). We also noted genes with a link to spermidine/putrescine transport (PP_0412-13), as well as a gene coding for a DUF2790 domain-containing protein. The latter was strongly downregulated when La supplemented cultures with those grown with Er or without Ln (log_2_FC -2.89, -2.62).

When looking at cluster 2, gene expression was elevated in all comparisons between cultures grown with La, Nd, or the Ln mix; and cultures supplemented with Er or no Ln. A weaker response was visible when Ln mix cultures were checked against, La, Nd, or Er. Cluster 2 comprised transcriptional regulator genes (PP_1683, PP_3756), genes encoding (heavy) metal sensor and transport proteins (PP_5139, PP_5383-84), a phasin gene (PP_5008), but also genes involved in Fe-S cluster assembly (PP_0843-46) (**Figure 5B**).

Cluster 3 included two copies of the chemotaxis-related adaptor protein CheW (PP_1489, PP_1491). It featured reduced expression in the two comparisons between La and Er, and between La and the control cultures, while expression was higher when contrasting Nd, and the Ln mix, against La. The genes belonging to cluster 4 included multiple TetR/Acr transcriptional regulator genes (PP_1961, PP_1968), as well as various ribosomal protein genes. We noted lower gene expression when comparing Nd and Ln mix cultures with Er and cultures without added Ln.

The addition of light/mixed Ln caused elevated gene expression of cluster 5 genes in comparison to the control set-up and cultures grown with Er. Genes for FeS-cluster assembly proteins were sensitive to Ln supplementation (log_2_FC -1.78 to 1.76), while the most responsive cluster 5 genes encoded metal-sensing, metal-binding proteins, and another DUF2790 domain-containing protein (PP_2969, PP_5732, PP_3494; log_2_FC -3.34 to 3.04). Cluster 6, covered in more detail in the next section, included lanthanome-related genes.

### Differential gene expression of lanthanome-associated genes

The lanthanome comprises all biomolecules, primarily proteins, directly involved in Ln utilization (28). In *P. alloputida* KT2440, Ln utilization is mediated through the *ped*-cluster (PP_2664 – PP_2680) that is linked to 2-phenylethanol uptake and conversion to phenylacetic acid (29). It includes *pedE*, *pedH*, as well as genes coding for the Ln-sensing two-component system PedS2/PedR2 (PP_2671/PP_2672), and an ABC transporter PedA1A2BC (PP_5538, PP_2669, PP_2668, PP_2667) facilitating cytoplasmic Ln uptake. Besides the *ped*-cluster, we also considered PQQ biosynthesis as part of the lanthanome. In addition, we looked at the pea-cluster (encoding genes of the 2-phenylethylamine pathway), since encoded proteins are also quinoprotein dehydrogenases.

Except for Er addition, the *ped*-cluster, including *pedE* and *pedH*, was downregulated upon Ln supplementation, (**Figure 6A**, **Table S4-13**) (log_2_FC values between -1.29 and -6.98). Only *pedE* (PP_2674) was differentially expressed when comparing incubations with Er and without added Ln (log_2_FC -0.96). Depending on the added Ln element, *pedE* gene expression differed by up to two orders of magnitudes (fold change of up to 126). Its highest gene expression was detected in samples without Ln (RPKM [reads per kilobase per million] 410.28 ± 56.55), followed by Er (RPKM 210.45 ± 27.53). Significantly lower gene expression was observed in samples supplied with light Ln or the Ln mix. Gene expression ranged between 4.45 ± 0.29 (Nd) and 3.25 ± 0.57 (La) RPKM. The gene expression ratio of *pedE* and *pedH* differed depending on the Ln supply (**Figure 6B**). In samples without Ln and with added Er, the *pedE*:*pedH* gene expression ratio was six and two, respectively. The *pedE*:*pedH* ratio shifted in favor of *pedH* in response to La, Nd, and the Ln mix. The expression of *pedH* was between 1.9-(Nd) and 4.2-fold (La) higher.

**Figure 6.**
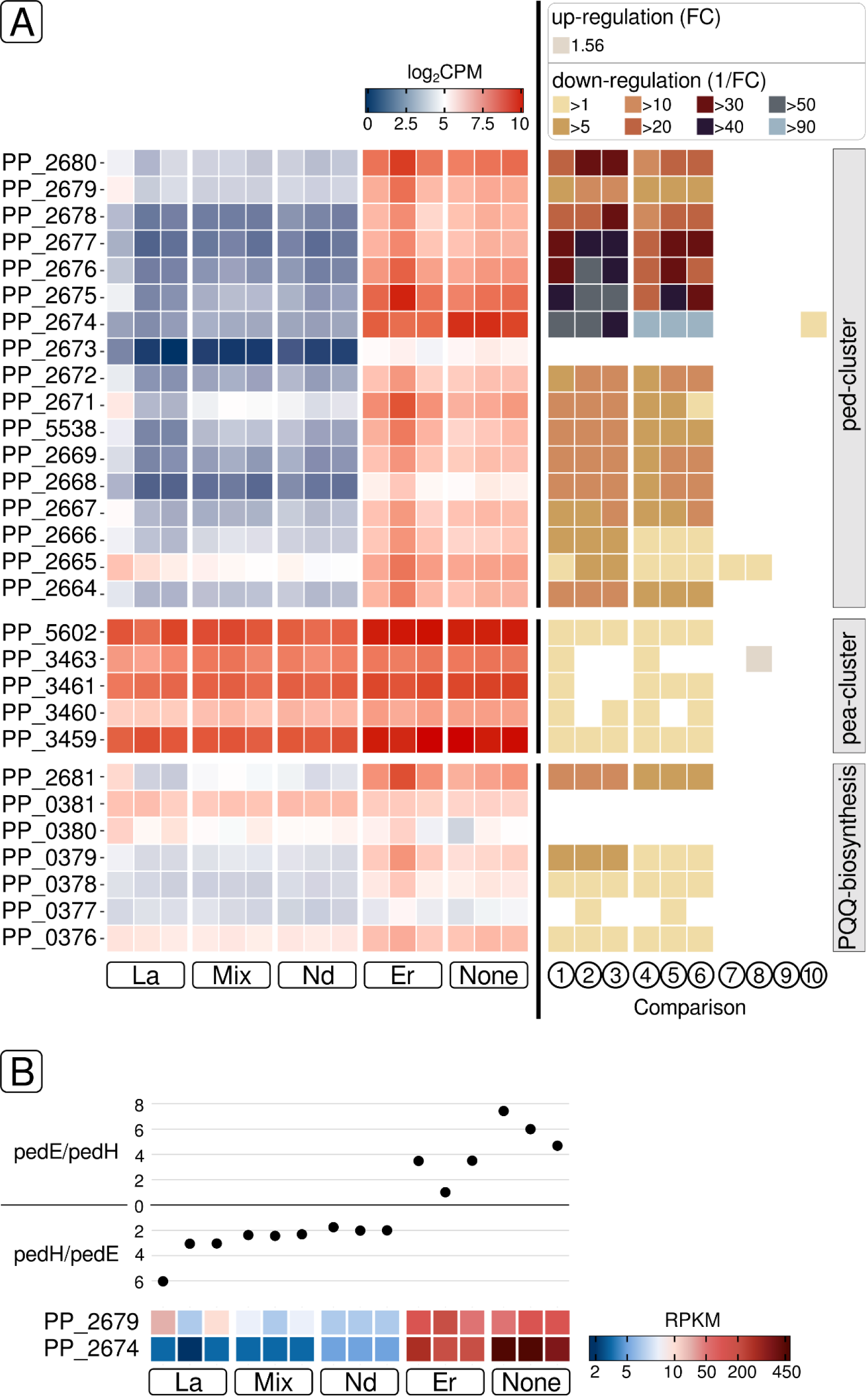
Groups of genes associated with the lanthanome were identified. They encode the *ped*-cluster, the *pea*-cluster, and genes involved in PQQ-biosynthesis. Gene expression is plotted based on log_2_CPM values (A, left panel). Changes in gene expression are shown for each comparison ((1) to (10) as described in Fig. 5)) based on the respective fold change (FC) or its reciprocal in case of down-regulation (A, right panel). The latter value (1/FC) describes the factor by which the expression of the respective gene is reduced. The Ln element-dependent gene expression ratios of ADH-encoding genes *pedE* (PP_2674) and *pedH* (PP_2679) were calculated based on RPKM values and plotted (B).

We noted a downregulation of PQQ biosynthesis-related genes (*pqqBCDE*; (PP_0379, PP_0378, PP_0376, PP_2681) when comparing La, Nd, and the Ln mix against Er and samples without Ln addition. Differential gene expression was not detected when light Ln samples were compared to each other or the Ln mix. There was no difference between the Er and no Ln samples. Genes associated with the *pea*-cluster (PP_3459 - PP_3461, PP_5602) were overall highly expressed (log_2_CPM between 6.21 ± 0.06 and 10.06 ± 0.09). Downregulation was observed when light Ln or the Ln mix were supplied (log_2_FC between -0.60 and -1.29).

## DISCUSSION

In *P. alloputida* KT2440, the Ln switch is rooted in the interplay between the Ca-dependent PQQ ADH PedE, and the Ln-dependent PQQ ADH PedH (21). PedH also functions as a sensory module and is, together with the PedS2R2 (PP_2671, PP_2672) two-component system (15), integral for Ln signaling. It was previously postulated that PedS2 phosphorylates its LuxR-type response regulator PedR2 in the absence of La, which in turn activates *pedE* and limits *pedH* expression (15). If La is available, PedS2 kinase activity is reduced, *pedE* expression is decreased, and *pedH* transcription is activated (15).

Our findings suggest that the Ln sensing and signaling machinery in *P. alloputida* KT2440 responds differently to utilizable and non-utilizable Ln. Utilizable Ln reduce Ln signaling gene expression. It is not clear if PedS2 binds heavier Ln, if heavier Ln reduce its activity, or if any other sensory elements are acting upstream of PedS2R2. The observed differences in gene expression between utilizable and non-utilizable for *pedE*, *pedH*, and *pedS2R2* indicate that Ln sensing and signaling in *P. alloputida* KT2440 are more complex than previously anticipated.

Our analyses suggest that the Ln switch is a dimmer switch. Compared to cultures without added Ln, we did not observe an upregulation of *pedH* in response to Ln supplementation. Except for Er, we noted the simultaneous downregulation of *pedE* and *pedH* in the presence of Ln, but also the reversion of their expression ratio. Previously carried-out promoter activity assays (21) showed an increase of *pedH* and a decrease of *pedE* promoter activity in the presence of La.

Our incubation experiments showed that the reduced and reversed expression ratio of *pedE*:*pedH* was no disadvantage. Modulating the pool of PQQ ADH depending on Ln availability might be a mechanism to optimize substrate utilization. PedH features a higher V_max_ and specific activity compared to PedE. At the same time, PedH has a much higher affinity for Ln than PedE for Ca (21). The downregulation of *pedE*, while *pedH* expression was unaltered when *P. alloputida* KT2440 was supplemented with Er supports the idea that non-utilizable Ln might interfere with PedSR2 activity and the modulation of the PQQ ADH pool. Enzymatic activity assays have shown that PedH activity is highest with LREM and absent with heavy Ln such as Er or Yb (21). Ln-dependent PQQ ADH can accommodate LREM and HREM with only minor changes in overall protein structure. However, differences in ionic radii and coordination number affect the catalytic efficiency of PQQ ADH and are potential reasons for bacteria preferring the bigger and more abundant LREM (30). Accidental mismetallation of PedH with non-utilizable Ln might explain the impaired growth observed with Er.

Droplet-based microfluidics combined with segmented flow is ideally suited for high-throughput cultivation and has been used for cultivating and characterizing various microbes, for instance, soil microorganisms such as *Rhodococcus* sp. and *Chromobacterium* (*31*), cyanobacteria (32), as well as plant stem cells (33). Our one-dimensional microfluidic screenings suggested higher fitness with increasing La concentration, which complied with the observed growth efficiency when using conventional cultivation. Taking into account that PedH is not active with Er (21), and the observed gene expression changes, the trend of higher final autofluorescence for Er concentrations above 75 nM did not reflect higher fitness. Instead, it might be an indicator of increased oxidative stress due to mismetallation (34, 35).

The performed two-dimensional screening suggested that the presence of the non-utilizable Er in low concentration had a positive effect on the final autofluorescence values if La was present in higher concentration. One potential explanation might be interference with the regulation of *pedE* and *pedH* gene expression. For now, the mechanism underlying this effect remains unanswered. In the future, coupling high-throughput microfluidic cultivation with downstream analyses, such as examining gene expression through RNAseq, might be a powerful strategy for studying Ln-related, phenotypic differences on multiple levels.

Observed growth patterns indicated that *P. alloputida* KT2440 interacts not only with light but also with heavy Ln. Intracellular Ln accumulation was previously shown for *M. extorquens* AM1 (18, 36) and Beijerinckiaceae bacterium RH AL1 (37). In the case of *M. extorquens* AM1, Ln are accumulated in the cytoplasm, while strain RH AL1 deposits Ln in the periplasm. In *M. extorquens* AM1 and Beijerinckiaceae bacterium RH AL1, Ln uptake into the peri- and cytoplasm is enabled by TonB-ABC transporter systems. The TonB-dependent receptor is in charge of uptake into the periplasm, and an ABC transporter facilitates translocation into the cytoplasm (18, 37, 38).

For *P. alloputida* KT2440 it was shown that cytoplasmic Ln uptake relies on the PedA1A2BC (PP_2267-69, PP_5538) ABC transporter (22). We could neither identify extra-nor intracellular Ln accumulations in *P. alloputida* KT2440 through electron microscopy. Single-cell ICP-MS (39) enabled us to quantify cell-associated Ln. Cell-associated Ln in Ln-utilizing microbes have so far not been analyzed on the single-cell level. Despite only one known Ln-dependent protein, PedH, the determined values of cell-associated Ln (0.0025 - 0.0068 % wet weight) are comparable to those known for Fe in *Escherichia coli* (40).

Previous studies addressing Ln-utilizing methylotrophs revealed that Ln supplementation affected primarily xox- and mxa-cluster genes (41–44). *M. aquaticum* 22A is the only example, for which the effect of light (La) and heavy Ln (Ho, Lu) on gene expression was investigated (42). Methylotrophy-related genes were found to be sensitive to La addition, while Ho and Lu had no noticeable effect on gene expression. We could show that Ln have a tremendous effect on gene expression in Beijerinckiaceae bacterium RH AL1, which does not possess Ca-dependent PQQ ADH and lacks the Ln switch. Up to 41% of the encoded genes were differentially expressed in strain RH AL1 dependent on Ln supplementation (45).

We observed a clear difference between utilizable and non-utilizable Ln regarding gene expression. The upregulation of ribosomal protein genes L10 and L7/12; involved in ribosome stalk formation, translation factor recruitment, and GTP hydrolysis (46); when comparing La with Er and no Ln supplementation, suggested an overall effect on protein biosynthesis. The differential expression of numerous transcriptional regulator genes hinted towards dedicated mechanisms for transmitting changes in Ln availability. Genes encoding DUF2790-containing proteins were among the most differentially expressed genes dependent on Ln supplementation. This family of proteins of unknown function is conserved within the Pseudomonadaceae. The genome of *P. alloputida* KT2440 includes seven genes coding for DUF2790-containing proteins, of which four were responsive to Ln supplementation, suggesting that this poorly characterized group of proteins might play a role regarding Ln utilization.

Some of the observed gene expression changes point towards Ln affecting the redox state of *P. alloputida* KT2440. Examples include the upregulation of genes linked to a spermidine/putrescine transport system when comparing La against Er and no Ln addition. Polyamines such as spermidine and putrescine function as antioxidants and signaling molecules (47, 48). When grown with La, Nd, or the Ln mix, genes associated with FeS cluster assembly were significantly more expressed, indicating higher demand, compared to cultures grown with Er or without Ln. Oxidative stress can cause an increased demand for functional FeS clusters (49). FeS cluster proteins are central to many forms of metabolism, including respiration, photosynthesis, as well as nitrogen fixation. FeS proteins can also function as sensors for intracellular oxygen and iron concentration (50).

Several genes for metal transporters showed differential gene expression when *P. alloputida* KT2440 was grown with Ln. Metalloproteins can accommodate different metals based on similar coordination geometry and the structure of the metal-binding site. Wehrmann and colleagues (22) could show that micromolar concentrations of Cu and Zn prevent the growth of a *P. alloputida* KT2440 *pedE* mutant when grown with nanomolar concentrations of Ln. Cu and Zn are known to strongly bind non-cognate metal-binding sites (51, 52).

*P. alloputida* KT2440 features two genes for phasin proteins (PP_5007, PP_5008), and one of them was significantly upregulated upon Ln supplementation. Scaffolds of phasin proteins surround polyhydroxybutyrate (PHB, a polyhydroxyalkanoate [PHA]) vacuoles (53). PHB vacuoles are not only linked to carbon storage. The presence of numerous proteins on their surface and their designation as carbonosomes highlights their assumed multifunctionality (54). PHB and related PHA are of economic relevance in the context of bioplastic production (55). For Beijerinckiaceae bacterium RH AL 1, we proposed that PHB plays a role in Ln uptake, periplasmic storage, and overall Ln homeostasis.

## CONCLUDING REMARKS

Taken together, we give evidence that Ln sensing in *P. alloputida* KT2440 is tuned towards utilizable Ln. Our findings support the idea that the Ln switch operates as a dimmer switch, modulating the pool of Ca- and Ln-dependent PQQ dependent on Ln availability. Ln-dependent gene expression changes have been opaque but were arguably skewed towards metal homeostasis and changes in the overall cellular redox state. The amount of cell-associated Ln raises the question if Ln are involved in processes other than VOC oxidation. The continued and in-depth study of Ln-utilization in organisms beyond methylotrophy is needed to fully unravel the relevance of Ln in microbial physiology and thus its potential impact on microbial ecology and potential biotechnological applications.

## MATERIALS AND METHODS

### Cultivation

*Pseudomonas alloputida* KT2440 (DSM #6125) was maintained on solid MP (*Methylobacterium* PIPES) medium (56) supplemented with 2-phenylethanol (5 mM) at 30 °C. Ln used for cultivation were supplemented as trichloride salts (retrieved in analytical grade from Carl Roth [Karlsruhe, Germany] and Sigma-Aldrich [Taufkirchen, Germany]). Incubation experiments were carried out with MP medium either in 96-well plates (BRANDplates®, Brand + CO. KG, Wertheim, Germany) (cultivation volume 300 µL), or acid-washed 150 mL Erlenmeyer flasks (cultivation volume 50 mL). Pre-cultures were grown with succinate as the carbon source (25 mM). More information is given in the **supplementary material**.

### Microfluidic cultivation

The setup used for microfluidic cultivation has been described in detail previously (57, 58) (**Figure S1**). We used a syringe pump with six dosing units and a six-port droplet generator from Cetoni GmbH (Korbußen, Germany) to generate droplets with different compositions. Details can be found in the **supplementary material**.

*Single-cell (sc) ICP-MS (inductively coupled plasma-mass spectrometry) analysis* Incubations were run as outlined before in 96-well plates, with three biological and six technical replicates. Incubations without Ln addition served as a control. Biomass was collected after the incubations reached OD_600nm_ values between 0.25 and 0.50. Technical replicates were pooled, harvested, and fixed with 500 µL of 2.5% [v/v] glutaraldehyde (in 100 mM cacodylate buffer) overnight at 4°C. Fixed and washed pellets were resuspended in 600 µL cacodylate buffer and cell numbers were determined using a counting chamber. scICP-MS measurements were performed using an 8900 inductively coupled plasma-mass spectrometer (ICP-MS/MS) (Agilent Technologies, Inc., Santa Clara, CA, USA) equipped with a high-efficiency sample introduction system (Glass Expansion, Port Melbourne, Australia) composed of a total consumption spray chamber and a microconcentric nebulizer (39). Data acquisition is further outlined in the **supplementary material**. For determining the wet weight portion of Ln, we took wet and dry weights of 1.7 g × L^-1^, and 0.4 g × L^-1^ per unit OD_600nm_ (59) as proxy. Assuming a cell number of 7.8 × 10^8^ × (mL × OD_600nm_)^-1^ (60), results in a wet weight per cell of approximately 2.2 pg.

### Electron microscopy

Transmission electron microscopy (TEM), and energy-dispersive X-ray spectroscopy (EDX) analyses were done as described previously (37, 45).

### RNA extraction, mRNA enrichment, and sequencing library preparation

Total RNA was extracted and mRNA enriched based on previously described methods (37, 38, 61). Sequencing libraries were prepared using the NEBNext Ultra II Directional RNA Library Prep Kit for Illumina (New England Biolabs, Germany). The size distribution of the libraries was checked by high-resolution gel electrophoresis with a Bioanalyzer instrument using the DNA 7500 Pico kit (Agilent Technologies). Libraries were quantified through fluorometry with a Qubit^TM^ fluorometer and dsDNA HS reagents (Thermo Fisher Scientific, Darmstadt, Germany).

### Sequencing and data pre-processing

An equimolar pool of the prepared libraries was sequenced with a NovaSeq 600 instrument (Illumina, San Diego, California, USA) in paired-end mode (2 × 100 bp). Sequencing was carried out by the sequencing core facility of the Leibniz Institute on Aging - Fritz Lipmann Institute (Jena, Germany). We assessed the quality of raw and trimmed sequences with *FastQC* (v0.11.9) (62). Adaptor- and quality-trimming (settings: minlen= 75, qtrim = rl, ktrim = rl, k = 25, mink= 11, trimq = 20) were carried out with *bbduk* (v38.26) (63) using its included database of common sequence contaminants and adapters. The filtering of rRNA-derived reads, the mapping of mRNA-derived reads, and the generation of the read count table for subsequent differential gene expression analysis are detailed in the **supplementary material**.

### Differential gene expression and cluster analysis

The R software framework for statistical computing (v4.2.1) and the package *edgeR* (v3.36.0) (64, 65) have been used to identify differentially expressed genes. We considered genes with changes in gene expression above |0.58| log_2_FC (fold change), gene expression values higher than 4 log_2_CPM (counts per million), and FDR values smaller than 0.05 for differential gene expression analysis. Data exploration including cluster analysis is explained in more depth in the **supplementary material**.

### Figure generation

We used the R software framework (v4.2.1) (66) for plotting and made use of the packages *ggplot2* (v3.3.6) (67), *gplots* (v3.1.3) (68), *ggpubr* (v0.4.0) (69), *cowplot* (v1.1.1) (70), and *upsetR* (v1.4.0) (71), including their respective dependencies. Figures have been finalized with inkscape (https://inkscape.org/).

### Data availability

RNAseq data sets are available via EBI/ENA ArrayExpress (accession: E-MTAB-13102) [https://www.ebi.ac.uk/arrayexpress/experiments/E-MTAB-13102/]). Details about data processing and analysis are also provided via the Open Science Framework (https://osf.io/ynsdc/). We also provide a reproducible, snakemake-based, (72, 73), workflow for RNAseq data processing (https://github.com/wegnerce/smk_rnaseq, release v0.1).

## ACKNOWLEDGEMENTS

The authors thank Josefine Bach, and Stefan Riedel (Friedrich Schiller University Jena, Aquatic Geomicrobiology Group) for help with cultivation work, especially growth monitoring. JC gratefully acknowledges the financial support provided by a habilitation scholarship from the Technische Universität Ilmenau. CEW thanks Lena Daumann and Sophie Gutenthaler-Tietze (Heinrich Heine University Düsseldorf) for helpful discussions while preparing the mansucript. This research was supported by the Deutsche Forschungsgemeinschaft (grant: WE6579/4-1, granted to CEW) and through the SFB 1127 ChemBioSys, project number 239748522.

## AUTHOR CONTRIBUTIONS

LG supervised SLB, designed and carried out experimental work, carried out RNAseq data analysis, and contributed to the first draft of the manuscript. SLB carried out cultivations, molecular work, and performed RNAseq data analysis. BN and JC established microfluidic cultivation for *P. alloputida* KT2440, carried out all microfluidic cultivations, and helped with interpreting data. JC contributed to the first draft of the manuscript. SH and TS adapted and established scICP-MS for *P. alloputida* KT2440, and collected and processed primary ICP-MS/MS data. MW acquired, processed, and analyzed EM data. JB carried out complementary experimental work. CEW designed the experimental work, carried out RNAseq data pre-processing and data analysis, acquired funding, and wrote the final version of the manuscript based on input from all co-authors.

